# Pandemic-scale phylogenetics

**DOI:** 10.1101/2021.12.03.470766

**Authors:** Cheng Ye, Bryan Thornlow, Alexander Kramer, Jakob McBroome, Angie Hinrichs, Russell Corbett-Detig, Yatish Turakhia

**Author notes:** Denotes equal contribution (Cheng Ye, Bryan Thornlow). **Disclaimer:** This paper was submitted for nomination to the Gordon Bell Special Prize for High Performance Computing-Based COVID-19 Research 2021. While our team was not selected as a finalist, we are posting this as pre-print since we think it could be of interest to the community. We expect modified and extended versions of this pre-print to appear in future publications.

## Abstract

Phylogenetics has been central to the genomic surveillance, epidemiology and contact tracing efforts during the COVD-19 pandemic. But the massive scale of genomic sequencing has rendered the pre-pandemic tools inadequate for comprehensive phylogenetic analyses. Here, we discuss the phylogenetic package that we developed to address the needs imposed by this pandemic. The package incorporates several pandemic-specific optimization and parallelization techniques and comprises four programs: UShER, matOptimize, RIPPLES and matUtils. Using high-performance computing, UShER and matOptimize maintain and refine daily a massive mutation-annotated phylogenetic tree consisting of all SARS-CoV-2 sequences available in online repositories. With UShER and RIPPLES, individual labs – even with modest compute resources – incorporate newly-sequenced SARS-CoV-2 genomes on this phylogeny and discover evidence for recombination in real-time. With matUtils, they rapidly query and visualize massive SARS-CoV-2 phylogenies. These tools have empowered scientists worldwide to study the SARS-CoV-2 evolution and transmission at an unprecedented scale, resolution and speed.

## 2. Justification

Our package:

- helps maintain possibly the largest-ever phylogenetic tree with millions of SARS-CoV-2 sequences, thus providing an unprecedented resolution for studying the pathogen’s evolutionary and transmission dynamics
- incorporates a new sequence on the phylogeny and sensitively identifies its recombination history within seconds, thereby enabling real-time monitoring of the virus

## 3. Performance attributes

| Performance Attribute | Our Submission |
| --- | --- |
| Category of achievement | peak performance, scalability |
| Type of method used | N/A |
| Results reported on the basis of | whole application including I/O |
| Precision reported | N/A |
| System scale | results measured on full-scale system |
| Measurement mechanism | timers |

## 4. Overview of the problem

As the SARS-CoV-2 virus spreads in the human population, it acquires new mutations. These mutations can render the virus more contagious, virulent or capable of evading the vaccines and antibody-based therapies (Campbell et al., 2021; Challen et al., 2021; Harvey et al., 2021). Therefore, from the early days of the pandemic, the global scientific community mobilized to monitor the viral mutations and the evolutionary dynamics with the help of genome sequencing (Lo and Jamrozy, 2020; Maxmen, 2021). The first SARS-CoV-2 genome sequence was deposited on an online database in January 2020 (Wu et al., 2020), and since then, over 4 million additional sequences have been shared through an extraordinary worldwide effort, with tens of thousands more being shared every day (Maxmen, 2021; McBroome et al., 2021). This vast volume of genomic data has provided invaluable insights into the evolution and spread of the virus, and has allowed public health officials and governments to respond to it in a timely fashion (Lam-Hine et al., 2021; Oude Munnink et al., 2020).

Phylogenetics has been a foundational tool in analyzing the genomic data for a public health response (Hodcroft et al., 2021). COVID-19 phylogenetics aims to infer the evolutionary relationships between the different SARS-CoV-2 genome sequences sampled from infected people and represent this information in the form of a tree, with individual sequences occupying the leaves of this tree (Lam et al., 2010). Sequences that are similar to each other tend to be grouped together in the tree, since they are likely to share a recent common ancestor. Phylogenetic trees play a crucial role in genomic surveillance, which involves tracking SARS-CoV-2 *variants* – each representing a different lineage in a phylogenetic tree – circulating in a given geographic region (da Silva Filipe et al., 2021; Deng et al., 2020), as well as for identifying and naming new variants (Rambaut et al., 2020). They also provide a powerful tool in establishing transmission links between seemingly unrelated infections, and in disambiguating community transmission from outside introductions (da Silva Filipe et al., 2021; Deng et al., 2020; Komissarov et al., 2021). Moreover, phylogenetic trees complement experimental techniques to identify the mutations that might have conferred increased transmissibility to the virus (Korber et al., 2020; Richard et al., 2021). Phylogenetic trees also find numerous other applications in epidemiology, including in estimating the reproduction number (R0) of the virus or its particular variant (Lai et al., 2020; Volz et al., 2021).

### 4.1 Phylogenetic placement

Many of these far-reaching phylogenetic applications require a comprehensive tree – without sub-sampling sequences – in order to unravel the true potential of the available genomic data. For example, transmission links with a sequence missing in the sub-sampled phylogenetic tree cannot be established. Likewise, sub-sampled phylogenetic trees could omit important lineages or sub-lineages corresponding to different variants. This can have adverse consequences on the downstream evolutionary and epidemiological studies. But maintaining a comprehensive phylogenetic tree of over 4 million available SARS-CoV-2 sequences, with tens of thousands of new sequences becoming available each day, is computationally prohibitive with pre-pandemic phylogenetic tools. We therefore developed UShER – an ultrafast, parallel *phylogenetic placement* tool that can maintain a comprehensive phylogenetic tree by incorporating new sequences (also referred to as samples) as they become available onto an existing phylogenetic tree (Turakhia, Thornlow, AS Hinrichs, et al., 2021).

### 4.2 Tree Optimization

The greedy strategy in phylogenetic placement of sequentially incorporating new sequences onto an existing tree can occasionally lead to a suboptimal tree structure. This can be mitigated by *tree optimization* programs, which use tree rearrangement to find a more optimal tree. Here too, previous programs are inadequate to handle the vast scale and speed of SARS-CoV-2 genome data. We therefore developed matOptimize – a low-memory, high-performance computing (HPC) software for fast optimization of trees based on pandemic-scale data. To date, UShER and matOptimize have managed to maintain and refine the phylogenetic tree consisting of all SARS-CoV-2 sequences.

### 4.3 Recombination detection

The COVID-19 pandemic has intensified the need for individual labs worldwide having primary access to sequencing data to respond rapidly to the emergence of new variants. One worrisome mechanism through which the virus can produce new variants is *recombination* (Simon-Loriere and Holmes, 2011). Through recombination, different variants of the virus that co-infect a host cell can exchange genetic material to form novel variants with drastic “jumps” in fitness (Burke, 1997; Simon-Loriere and Holmes, 2011). Detecting recombination from viral genomes is so computationally demanding that prior tools could only handle up to a few thousand viral sequences – a far cry from the scale of millions that is needed to study SARS-CoV-2 recombination comprehensively. We show here how our tools UShER and RIPPLES (Turakhia, Thornlow, A Hinrichs, et al., 2021) empower individual research labs to provide a real-time response, not only in incorporating their sequences onto a global phylogeny, but also in uncovering evidence for recombination from a massive search space. RIPPLES can also be used in an HPC setting to comprehensively detect recombination events from the entire SARS-CoV-2 phylogeny within a few hours.

## 5. Current state of the art

### 5.1 Phylogenetic placement

In the recent decades, phylogenetic tools have been hugely focused on probabilistic approaches, namely maximum likelihood (ML) and Bayesian inference (BI). These approaches are guided by a probabilistic model of evolution (Durbin, 1998). For comparative genomics or molecular epidemiology of rapidly evolving pathogens, ML and BI techniques are indeed more accurate than the simplistic, model-free approaches of maximum parsimony (MP) or distance-based clustering (Page and Holmes, 1998). This is because when the sequence divergence is high, model-free approaches are more susceptible to *long branch attraction* (Bergsten, 2005; Felsenstein, 1978). State-of-the-art phylogenetic placement tools, including EPA-NG (Barbera et al., 2019), PPLACER (Matsen et al., 2010) and IQ-TREE2 (Minh et al., 2020), are all based on maximum likelihood. Each tool accepts (i) a reference tree and (ii) a FASTA-formatted file containing the multiple-sequence alignment (MSA) of all sequences, including the sequences to be placed on the reference tree. Placement typically starts with a pre-placement phase that uses fast heuristics to select promising candidate branches in the reference tree for each query sequence, followed by a detailed placement phase which evaluates the likelihood scores for candidates in detail through a thorough numerical optimization (Barbera et al., 2019). Ample parallelism available in the form of independent candidate branches is exploited. Despite highly-efficient implementations, ML-based placement tools have a high compute and memory requirement and cannot scale beyond a few tens of thousands of sequences – making them highly inadequate in the context of the COVID-19 pandemic. Recently, distance-based placement algorithms are also being explored (Balaban et al., 2020).

### 5.2 Tree optimization

Unlike phylogenetic placement, tree optimization has been widely studied for both maximum likelihood (Minh et al., 2020; Price et al., 2010) and maximum parsimony (Goloboff and Catalano, 2016). TNT (Goloboff and Catalano, 2016) is the most efficient parsimony-based tree optimization tool and scales well beyond the limits of likelihood-based tools. TNT (i) takes as input an existing tree in Newick format and the MSA of its sequences in FASTA format, (ii) computes the parsimonious state assignments for every node of the tree for every alignment site, and (iii) uses tree rearrangement in the form of tree bisection and reconnection (TBR) to improve parsimony score. TNT keeps multiple equally parsimonious candidate trees and uses tree drifting during the optimization to avoid getting stuck in local optima. TNT also provides several heuristics, such as ratchet search, to speed up the search (Goloboff, 1999). We found the *sectorial search* mode in TNT (Goloboff, 1999), which splits the original tree into subtrees to be optimized independently and merged back, to be the most computationally efficient heuristic for optimizing the SARS-CoV-2 phylogenies. TNT does not provide a multithreaded implementation, but we were able to adapt its scripts (see Section 7) to parallelize the optimization over several processes.

### 5.3 Recombination detection

Viral recombination has been studied widely in the past for a variety of viruses, including coronaviruses (Anthony et al., 2017; Hon et al., 2008; Koelle et al., 2017). Recombination Detection Program (RDP) is a widely-used program for recombination detection in viruses and provides implementation for a number of well-studied techniques in the literature (Martin et al., 2015; Martin and Rybicki, 2000). Broadly, RDP accepts a multiple alignment of sequences and scans each triplet of sequences in the alignment to discover whether one sequence of the triplet (viz., the putative recombinant) is a mosaic of the other two (viz., the parents of the recombinant). Specifically, RDP finds the recombination-informative signal in the alignment, i.e. sites in which two sequences share an allele that is different in the third, and then uses a number of heuristics to infer the recombination breakpoints that would best explain the signal. Because of the complexity of its search space, RDP analysis is limited to a few thousand viral sequences (Martin et al., 2015: 4).

## 6. Innovations realized

Our phylogenetic package achieves orders of magnitude speedup over the prior art through several domain-specific optimizations in the algorithms and data structures and efficient implementation of our parallel algorithms. Figure 1 illustrates these innovations.

**Figure 1:**
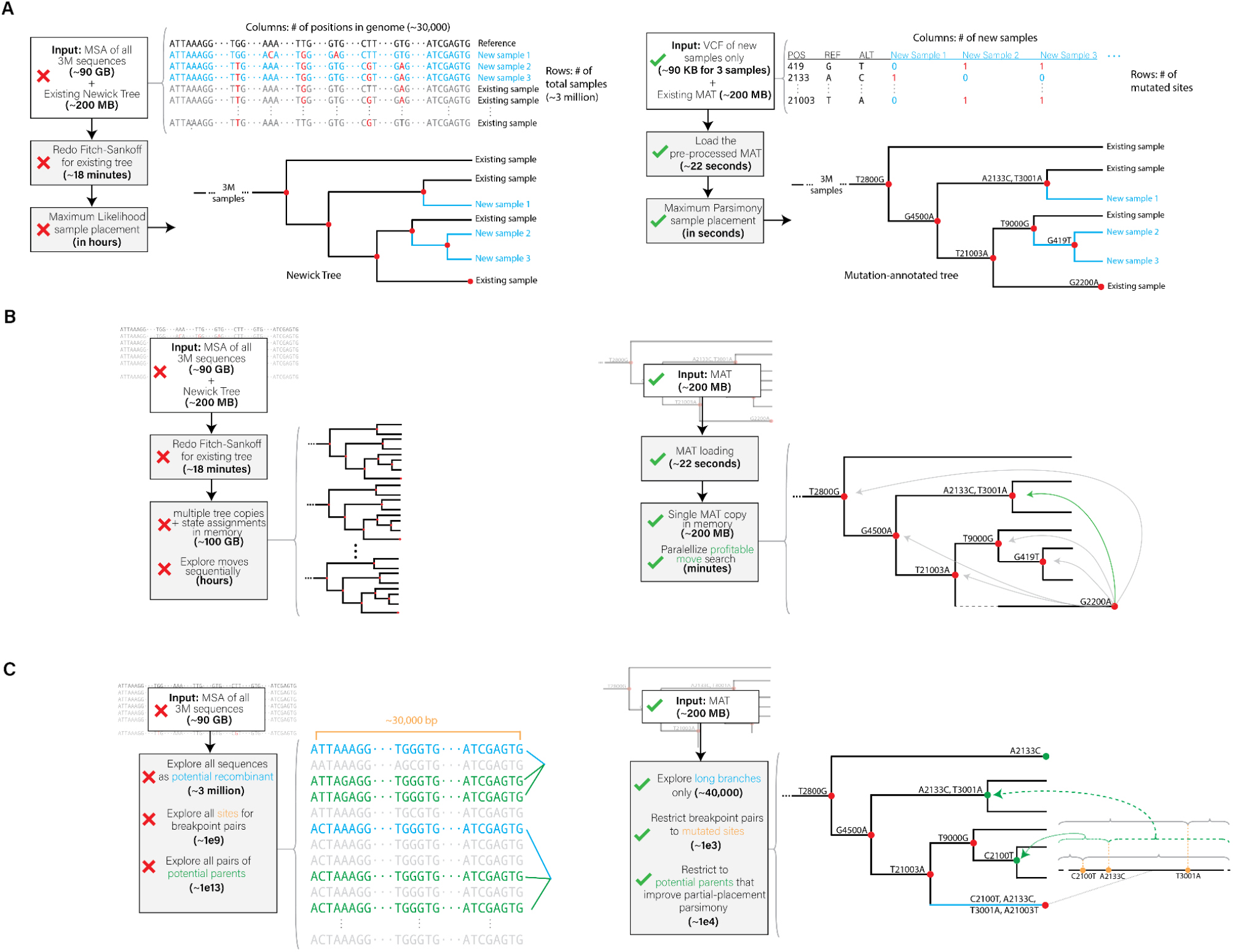
Innovative optimizations realized in (A) UShER, (B) matOptimize and (C) RIPPLES for phylogenetic placement, tree optimization and recombination detection, respectively. The left side shows a representative illustration of the prior approaches and the right side illustrates the approach used in our tools.

### 6.1 Algorithmic innovations

Pandemic-specific algorithmic innovations form the basis of our phylogenetic package efficiency. For example, we observed that in SARS-CoV-2 phylogenetics, long branch attraction is not an issue – a newly-sequenced SARS-CoV-2 genome is separated by less than one mutation, on average, from its closest neighbor in the global phylogeny. Unsurprisingly, we found ML and MP trees to be practically consistent, but the MP trees had better interpretability since individual mutations could be confidently labelled on internal branches, as done in our package using a novel data object called mutation-annotated tree (MAT) (Turakhia, Thornlow, AS Hinrichs, et al., 2021). MAT is a protocol buffer based file format as well as an internal data structure used in our package (Turakhia, Thornlow, AS Hinrichs, et al., 2021). Our phylogenetic package leverages the MAT and MP algorithms to avoid the expensive likelihood computations in probabilistic approaches.

A key algorithmic innovation that the MAT format enables is *pre-processing*, which is critical for the speedup of phylogenetic placement in UShER. MAT can be stored as a space-efficient pre-processed object (see Section 6.2), obviating the need to load the bulky sequence or variation data files for existing samples during each execution. It also avoids the need to recompute the parsimony state assignments internal to the tree using the computationally-intensive, dynamic programming-based Fitch-Sankoff algorithm (Fitch, 1971; Sankoff, 1975). The parallel Fitch-Sankoff algorithm using 20 CPU threads takes 18 minutes for a 3 million sample tree, approximately 50x slower than simply loading its MAT. Once the MAT is loaded and the new samples incorporated, the resulting MAT data structure could be stored to an output file within seconds to allow another round of placement over and above the current round. Besides pre-processing, UShER uses several additional algorithmic optimizations to speed up the placement, including condensing identical sequences into a single node and early termination of the symmetric set difference computation. These techniques are described in more detail in our previous publication (Turakhia, Thornlow, A Hinrichs, et al., 2021).

Our package also makes extensive use of *search space pruning* based on phylogenomic insights. At first, it may seem that RIPPLES has an intractable search space. An arbitrary choice of recombinant sequence, two parent sequences and two breakpoints would result in a search space that is cubic to the number of sequences – in millions – and quadratic to the sequence length – in tens of thousands. RIPPLES leverages the insight that a detectable recombinant sequence must have a long branch in its phylogenetic ancestry to greatly reduce the search space. RIPPLES also restricts the breakpoints to mutated sites in the recombinant and parent sequences. Parents are restricted to a small number of nodes – typically in tens – that reduce the parsimony score of partial phylogenetic placement (i.e. on ignoring sites outside the breakpoint regions) above an appropriate minimum threshold for detectable recombination.

### 6.2 Efficient data structures

The mutation-annotated tree (MAT) is a highly efficient data structure and key performance driver in our tools. Previous phylogenetic packages have typically relied on FASTA or VCF formats to store sequence and variation data. An uncompressed MAT file of 3 million SARS-CoV-2 sequence requires only 136 MB to encode basically the same information that is contained in an 88 GB FASTA or 174 GB VCF file. With MAT, matOptimize can start optimizing this tree after a small 22-second delay of loading the MAT file. In comparison, TNT can spend hours to load the alignment file and compute the internal state assignments before starting optimization (see Results, Figure 3B). Likewise, with mutations pre-annotated on the phylogenetic tree, RIPPLES can avoid scanning through the entire alignment file and focus solely on mutated sites.

### 6.3 Parallel implementation

We designed the algorithms to be embarrassingly parallel with limited synchronization overheads. Our package uses Intel’s TBB library (https://github.com/oneapi-src/oneTBB) to take advantage of available parallelism in multicore processors. In UShER, the placement search is parallelized over millions of internal nodes and leaves of the tree where a new sample could be placed. In matOptimize, tree arrangement is explored starting from all possible source nodes, in millions. This contrasts TNT, where each process optimizes the entire tree, and broadcasts the best tree to all other processes. Since the tree object remains immutable throughout the search phase in matOptimize, each thread can find profitable rearrangements independently, with minimal inter-thread communication. In RIPPLES, the partial phylogenetic placement of individual segments, as well as the parent and breakpoint search, is parallelized as well.

We also parallelized our tools for use in a CPU cluster. For parallelizing UShER over CPU nodes, we had to implement a *merge* operation in matUtils (McBroome et al., 2021). Briefly, this operation accepts two input MATs, checks if the subtrees resulting from common samples are consistent (i.e. have the same topology and mutation annotations), and then places the remaining samples in the common subtree using placement information derived from input MATs to narrowly restrict the search. This allows new samples to be independently placed on a MAT using UShER and the resulting MATs can then be rapidly merged together using a parallel reduction tree of matUtils merge to produce the final MAT (Figure 2). In matOptimize, the search for profitable moves is parallelized across nodes using MPI (https://www.mpich.org/) during iterations. An iteration begins with the main process distributing source nodes to worker processes. Then, the worker processes evaluate all possible subtree pruning and grafting (SPR) moves (without applying moves) from the assigned source nodes and periodically return the profitable moves to the main process, which applies those moves to update the tree and begin the next iteration. For RIPPLES, we implemented an option to restrict the search to a specific range of potentially recombinant nodes that allowed us to trivially parallelize it over the cluster without any communication or synchronization requirements.

**Figure 2:**
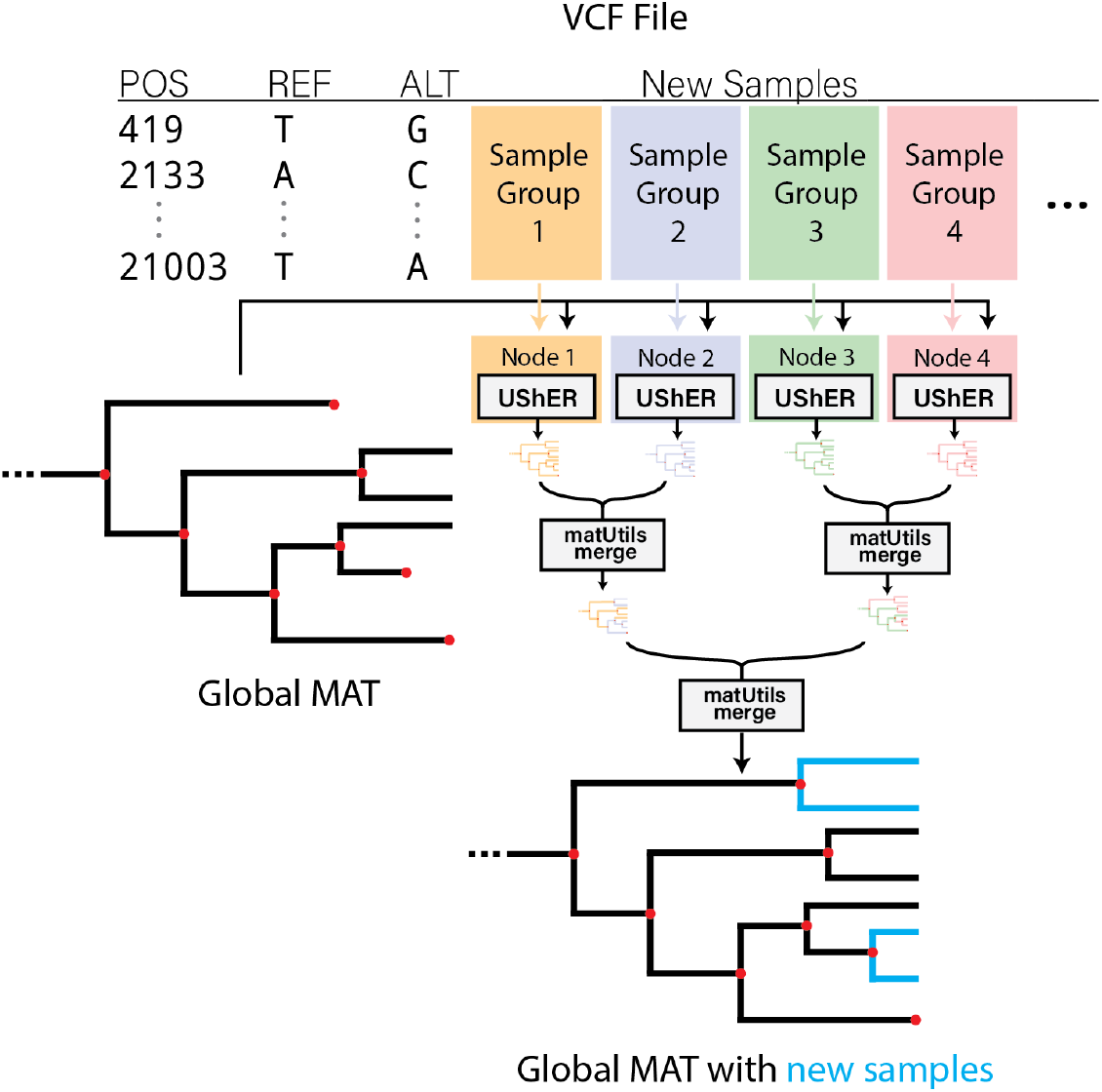
For parallelizing phylogenetic placement over multiple CPU nodes, we split up the VCF containing new samples uniformly and distribute them over independent CPU nodes, each executing UShER to place the corresponding samples on the base mutation-annotated tree (MAT). The resulting MAT files are then merged using a parallel reduction tree of matUtils merge into a single output MAT containing new samples.

## 7. How performance was measured

### 7.1 Reference tree and sequence data

We collected SARS-CoV-2 genome sequence data from major online databases: GISAID (Shu and McCauley, 2017), COG-UK (Nicholls et al., 2020), GenBank (Clark et al., 2016) and CNCB (https://bigd.big.ac.cn/ncov/release_genome), and used it to build a comprehensive SARS-CoV-2 phylogeny based on the methodology described in (McBroome et al., 2021). We used the GenBank MN908947.3 (RefSeq NC_045512.2) sequence as the reference for rooting the tree, as well as for calling variants in individual samples. We used mafft (version v.7.471) (Katoh and Standley, 2013) for aligning the sequences and faToVcf utility included in the UShER package to produce the required VCF files for our analyses. For our experiments, we used the sampling date metadata to derive from our comprehensive tree two subtrees containing the earliest 100K and 1M samples, referred to as 100K-sample tree and 1M-sample tree, respectively.

### 7.2 Experimental setup and comparison baseline

All experiments were performed on the Google Cloud Platform (GCP) and are easily reproducible. We used a recent version (commit e83dc5) of our package available at https://github.com/yatisht/usher for comparison against state-of-the-art tools with similar functionality as well as scaling analysis. Since our phylogenetic package is memory-efficient, we could use CPU-optimized E2 instances for our package but were required to use memory-optimized instances for some competing tools. In such cases, we opted for iso-cost comparison such that the hourly cost was roughly the same for both instances.

For phylogenetic placement, we compared UShER with IQ-TREE2 (version 2.1.3). Since IQ-TREE2’s memory requirement is prohibitively high for the 1M-sample tree, we compared the performance of placing 1000 new SARS-CoV-2 samples on the 100K-sample tree using the command options `-n 0 -m JC --suppress-list-of-sequences -nt 16`. We used the e2-highmem-16 instance (16 vCPUs, 128 GB, $0.72/hour) for IQ-TREE2 and e2-highcpu-32 instance (32 vCPUs, 32 GB, $0.79/hour) for UShER. Both programs are multithreaded and used all available vCPUs on the instance.

For tree optimization, we compared matOptimize with TNT (June 2021) using the 1M-sample tree. In particular, we used the sectorial search (Goloboff, 1999) in TNT, which we found to be the most efficient option for optimizing SARS-CoV-2 phylogeny, using the script at https://github.com/yceh/matOptimize-experiments/blob/master/tnt_parallel_sect.run. TNT is not multithreaded, but we managed to adapt the combosearch script (http://www.lillo.org.ar/phylogeny/tnt/scripts/combosearch.run) to parallelize it over several processes. We used m1-ultramem-40 instance (40 vCPUs, 961 GB, $6.30/hour) for TNT, with 8 processes, which we calculated to be the upper limit for the available memory on the instance. We used 7 e2-highcpu-32 instances for matOptimize that cumulatively have an hourly cost ($5.54/hour) comparable to the m1-ultramem-40 instance.

For recombination detection, we compared RIPPLES with OpenRDP (https://github.com/PoonLab/OpenRDP), which is a Python adaptation of RDP5 (Martin et al., 2021), using the n2d-highcpu-224 instance. Due to prohibitively long runtimes of OpenRDP, we could only compare the two tools on a small dataset of 1000 SARS-CoV-2 sequences for the typical pandemic scenario of determining whether a new sequence is a recombinant of the previous sequences. Our recombinant sequences were based on the compelling evidence provided in (Jackson et al., 2021). We used these sequences to derive the 1000 SARS-CoV-2 sequences using `matUtils extract -k 50` for each node in each trio representing each instance of recombination found in our global tree that was ancestral to the samples cited in (Jackson et al., 2021). Since OpenRDP is not multithreaded, we parallelized it by creating 498,501 OpenRDP jobs (using options `-m openrdp -rdp`) for each sequence triplet formed by combining the new sequence with every unique pair of sequences (one for each parent) from the 1000 SARS-CoV-2 sequences. We ran 224 jobs, one per vCPU, in parallel using GNU parallel utility (Tange, 2011). Because RIPPLES requires the new sequence to be already placed in the phylogenetic tree, we included the time to place the sample on the existing tree using UShER in the RIPPLES runtime for an approximately level comparison.

We also measured the peak memory usage of all programs using the proc filesystem (https://man7.org/linux/man-pages/man5/proc.5.html) in Linux.

### 7.3 Scaling analysis

We performed strong and weak scaling analysis for UShER, matOptimize and RIPPLES using the 1M-sample tree and e2-highcpu-32 instances, varying the number of instances from 2 to 32. For UShER and RIPPLES, parallel GCP instances were launched using the dsub utility (https://github.com/databiosphere/dsub). For UShER, scaling analysis was performed for placing 100K new samples on the reference tree. The time for parallel reduction merge using matUtils was included in the analysis and intermediate MAT files between the different stages of the reduction tree were communicated between instances through the Google Cloud Store (GCS) using the gsutil tool (https://cloud.google.com/storage/docs/gsutil), with the dependent instance waiting in a loop for its input MAT file to become available. RIPPLES was trivially parallelized through a static partitioning of the 9391 branches with 3 or more mutations and 5 or more mutations in the tree that were explored for a recombination event. For weak scaling, the problem size was also defined in terms of the number of long branches explored. For matOptimize, MPI (mpich 3.4.3; https://www.mpich.org/) was used for parallelizing search over multiple instances. A 1 million sample tree was optimized for 5 iterations with radius set to 1024, which amounts to SPR moves spanning the whole tree. Problem size was defined in terms of the number of source nodes explored for subtree pruning and regrafting (SPR) moves and for weak scaling, source nodes to search were randomized and divided uniformly between instances at each iteration.

## 8. Performance Results

### 8.1 Speedup analysis

Figure 3 highlights the orders of magnitude improvement in runtime and a large factor improvement in peak memory that our phylogenetic package achieves relative to state-of-the-art tools. UShER placed 1000 new samples on the 100K-sample tree in just 15.4 seconds using 92 MB of RAM, achieving 1439-fold speedup and 1300-fold improved memory-efficiency compared to IQ-TREE2. On tree optimization, TNT spent over 8 hours in just loading the alignment and computing the parsimony states before beginning optimization. However, in just over one hour, matOptimize completed its optimization. Even after 24 hours (a limit we imposed for daily optimization required in the pandemic) of TNT execution, the matOptimize tree remained more parsimony-optimal. For recombination detection, placing a new sample on the 1K-tree using UShER and flagging it as a recombinant using RIPPLES took a fraction of a second, but required over an hour using OpenRDP. RIPPLES is also 3 times more memory-efficient and as sensitive as OpenRDP for this dataset. On the 1M-sample tree, a new sample could be placed using UShER and inferred for recombination using RIPPLES in 35.65 seconds on average, which enables real-time monitoring of the virus for recombination.

**Figure 3:**
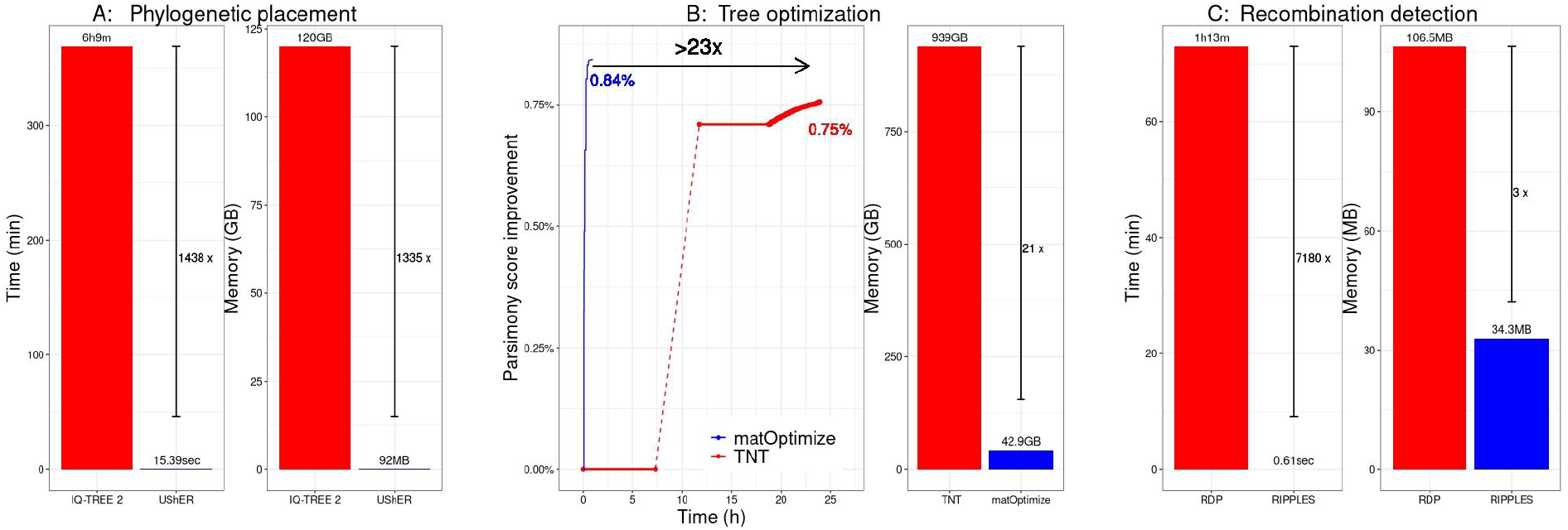
Comparison of our phylogenetic package with previous state-of-the-art tools for (**A**) phylogenetic placement, (**B**) tree optimization and (**C**) recombination detection. Our tools achieve large improvements in runtime (left) as well as peak memory requirements (right).

### 8.2 Strong scaling analysis

Figure 4 shows the strong scaling analysis of UShER, matOptimize and RIPPLES.

**Figure 4:**
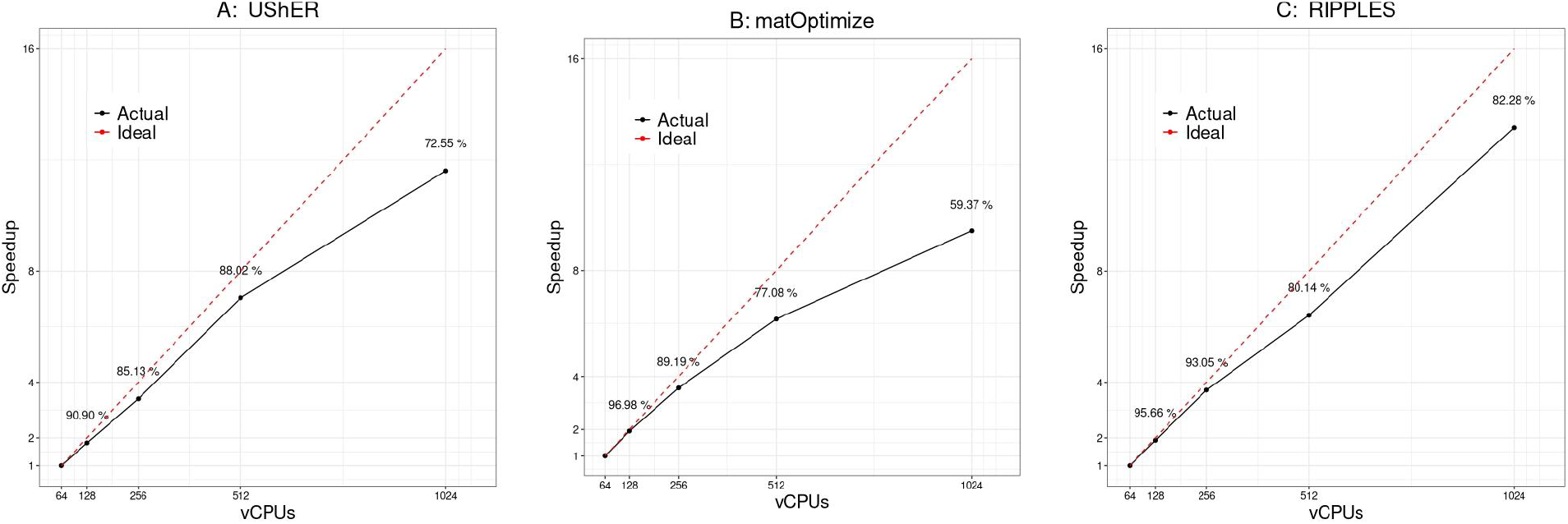
Strong scaling analysis for (**A**) UShER, (**B**) matOptimize and (**C**) RIPPLES.

UShER, in placing 100K new samples on the 1M-sample tree, maintains a strong scaling efficiency of over 85% until 512 vCPUs are used, after which point it drops to 72.6% at 1024 vCPUs (Figure 4A). Figure 5 illustrates the major factor behind UShER’s lost efficiency at high parallelism. The number of parallel reduction steps required for merging MATs increases with parallelism (Figure 2) and starts dominating the runtime at higher levels of parallelism. At 1024-way parallelism, the merge phase requires over 8 minutes and constitutes 21.7% of the total runtime, compared to only 1% at 64-way parallelism. Faster heuristics for matUtils merge could ameliorate this issue, which we plan to explore in the future.

**Figure 5:**
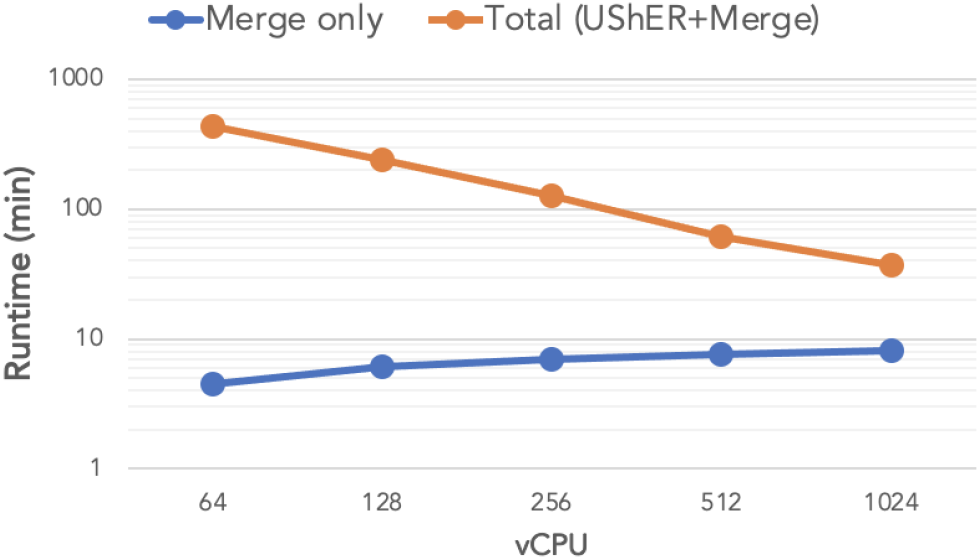
Total time (in orange) to place 100K new samples on the 1M-sample tree using UShER followed by parallel reduction using matUtils merge, with the merge component shown separately (in blue), for different levels of parallelism.

**Figure 6:**
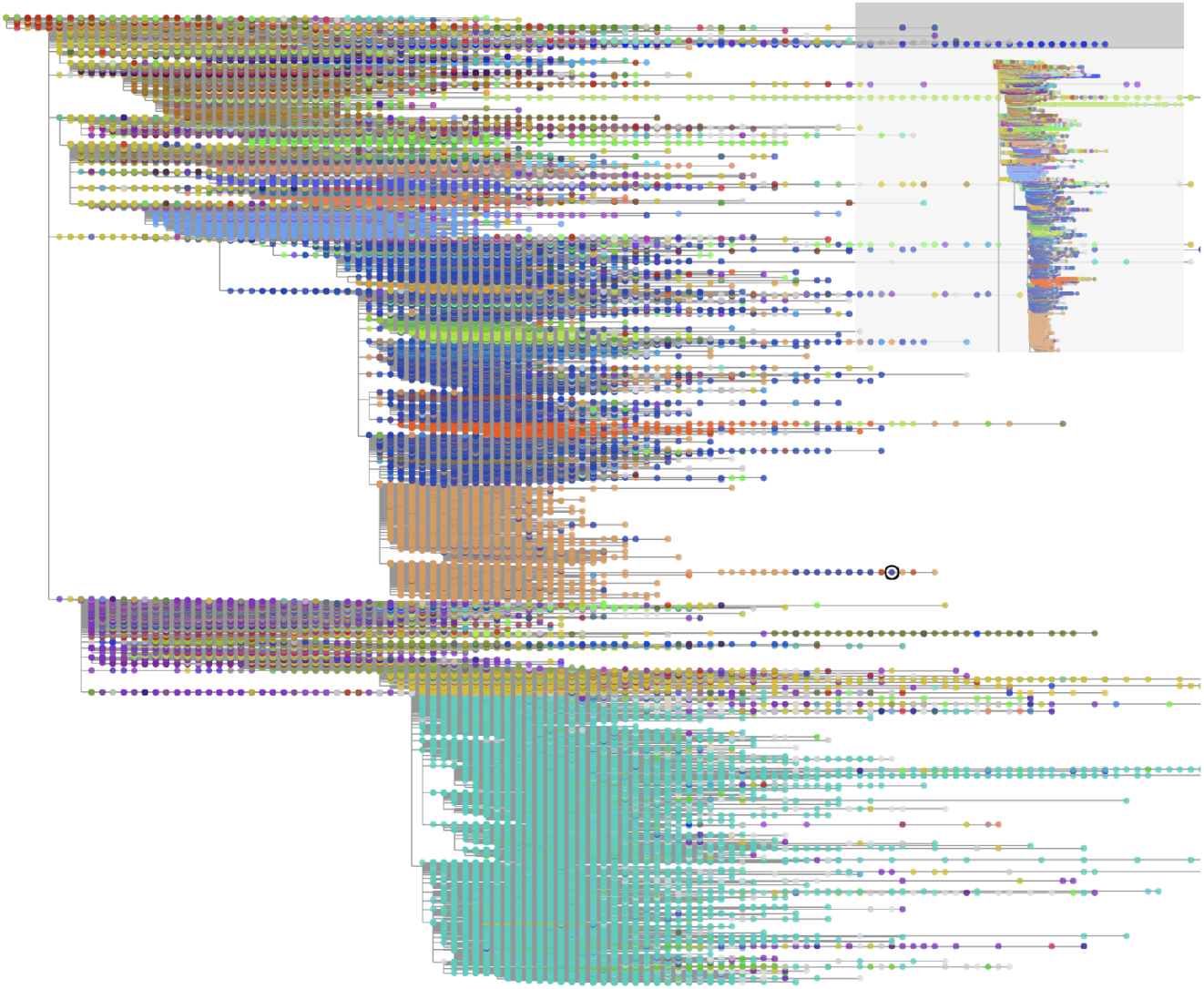
A comprehensive SARS-CoV-2 phylogeny as of October 3, 2021, containing 4,111,711 samples. The tree was built with the help of UShER and matOptimize and is visualized using CoV2Tree tool (https://cov2tree.org/). Each dot corresponds to a leaf, representing a single SARS-CoV-2 sample colored based on its Pango lineage assignment (https://pangolin.cog-uk.io/).

For matOptimize, we found a rapid deterioration of strong scaling efficiency with parallelism (Figure 4B). This is because through our highly efficient implementation of the union construct and incremental update methods (Goloboff, 1996), the parallel search for profitable moves was very fast for the 1M-sample tree relative to the sequential step of applying the moves, For example, with 1024 vCPUs, the entire matOptimize run required only 11.5 minutes, with the parallel search phase requiring less than 1.5 minutes on each iteration (7.5 minutes in total). We expect strong scaling efficiency to improve as the tree gets bigger.

RIPPLES achieved a strong scaling efficiency of over 80% for comprehensively detecting recombinants from the 1M-sample tree at all parallelism levels (Figure 4C). Parallel slack is the main factor behind lost efficiency at high parallelism – some long branches are 10-100 fold slower to analyze than the average and result in long tail latency jobs. Dynamic load balancing could improve the scaling efficiency of RIPPLES. Additionally, we also observed that some slack was created by the difference in time to spin up the GCP instances using dsub – some instances took over a minute longer to load than the rest.

### 8.3 Weak scaling analysis

Table 1 provides the weak scaling analysis for UShER, matOptimize and RIPPLES. All tools have a weak scaling efficiency above 70% and the causes behind the lost efficiency are the same as discussed in Section 8.2. The non-deterministic nature of our algorithms causes some non-monotonic variation of parallel efficiency in some cases.

**Table 1:** Weak scaling analysis for (**A**) UShER, (**B**) matOptimize and (**C**) RIPPLES.

| A. UShER |  |  |
| --- | --- | --- |
| vCPU | Samples placed | Time |
| 64 | 6.25K | 26m 48s |
| 128 | 12.5K | 28m 22s |
| 256 | 25K | 30m 41s |
| 512 | 50K | 33m 36s |
| 1024 | 100K | 37m 07s |
| B. matOptimize |  |  |
| vCPU | Source nodes explored | Time |
| 64 | 39789 | 10m 45s |
| 128 | 79577 | 11m 54s |
| 256 | 159154 | 11m 51s |
| 512 | 318308 | 11m 58s |
| 1024 | 636616 | 11m 30s |
| C. RIPPLES |  |  |
| vCPU | Long branches explored | Time |
| 64 | 587 | 49m 29s |
| 128 | 1174 | 53m 33s |
| 256 | 2348 | 54m 33s |
| 512 | 4696 | 56m 18s |
| 1024 | 9391 | 55m 52s |

## 9. Acknowledgments

We thank Devika Torvi for her contributions to matUtils merge. We thank Sneha D. Goenka for helpful tips on using Google Cloud Platform. We thank Willi Henning Society, for the TNT software.

This work was supported by funding from the National Institutes of Health (awards T32HG008345 and F31HG010584), the Centers for Disease Control and Prevention (BAA 200-2021-11554), University of California Office of the President Emergency COVID-19 Research Seed Funding Grant (award R00RG2456), Alfred P. Sloan Foundation and generous donations by the Eric and Wendy Schmidt Foundation.

## References

Anthony SJ, Gilardi K, Menachery VD, et al. (2017) Further Evidence for Bats as the Evolutionary Source of Middle East Respiratory Syndrome Coronavirus. mBio Schultz-Cherry S (ed.) 8(2). DOI: 10.1128/mBio.00373-17.

Balaban M, Sarmashghi S and Mirarab S (2020) APPLES: Scalable Distance-Based Phylogenetic Placement with or without Alignments. Systematic Biology 69(3): 566–578. DOI: 10.1093/sysbio/syz063.

Barbera P, Kozlov AM, Czech L, et al. (2019) EPA-ng: Massively Parallel Evolutionary Placement of Genetic Sequences. Systematic Biology 68(2): 365–369. DOI: 10.1093/sysbio/syy054.

Bergsten J (2005) A review of long-branch attraction. Cladistics 21(2): 163–193. DOI: 10.1111/j.1096-0031.2005.00059.x.

Burke DS (1997) Recombination in HIV: an important viral evolutionary strategy. Emerging Infectious Diseases 3(3): 253–259.

Campbell F, Archer B, Laurenson-Schafer H, et al. (2021) Increased transmissibility and global spread of SARS-CoV-2 variants of concern as at June 2021. Eurosurveillance 26(24). DOI: 10.2807/1560-7917.ES.2021.26.24.2100509.

Challen R, Brooks-Pollock E, Read JM, et al. (2021) Risk of mortality in patients infected with SARS-CoV-2 variant of concern 202012/1: matched cohort study. BMJ 372. British Medical Journal Publishing Group: n579. DOI: 10.1136/bmj.n579.

Clark K, Karsch-Mizrachi I, Lipman DJ, et al. (2016) GenBank. Nucleic Acids Research 44(D1): D67–72. DOI: 10.1093/nar/gkv1276.

da Silva Filipe A, Shepherd JG, Williams T, et al. (2021) Genomic epidemiology reveals multiple introductions of SARS-CoV-2 from mainland Europe into Scotland. Nature Microbiology 6(1). 1. Nature Publishing Group: 112–122. DOI: 10.1038/s41564-020-00838-z.

Deng X, Gu W, Federman S, et al. (2020) Genomic surveillance reveals multiple introductions of SARS-CoV-2 into Northern California. Science 369(6503). American Association for the Advancement of Science: 582–587. DOI: 10.1126/science.abb9263.

Durbin R (ed.) (1998) Biological Sequence Analysis: Probabalistic Models of Proteins and Nucleic Acids. Cambridge, UK: New York: Cambridge University Press.

Felsenstein J (1978) Cases in which Parsimony or Compatibility Methods will be Positively Misleading. Systematic Biology 27(4): 401–410. DOI: 10.1093/sysbio/27.4.401.

Fitch WM (1971) Toward Defining the Course of Evolution: Minimum Change for a Specific Tree Topology. Systematic Biology 20(4). Oxford Academic: 406–416. DOI: 10.1093/sysbio/20.4.406.

Goloboff PA (1996) Methods for Faster Parsimony Analysis. Cladistics 12(3): 199–220. DOI: 10.1111/j.1096-0031.1996.tb00009.x.

Goloboff PA (1999) Analyzing Large Data Sets in Reasonable Times: Solutions for Composite Optima. Cladistics 15(4): 415–428. DOI: 10.1111/j.1096-0031.1999.tb00278.x.

Goloboff PA and Catalano SA (2016) TNT version 1.5, including a full implementation of phylogenetic morphometrics. Cladistics 32(3): 221–238. DOI: 10.1111/cla.12160.

Harvey WT, Carabelli AM, Jackson B, et al. (2021) SARS-CoV-2 variants, spike mutations and immune escape. Nature Reviews Microbiology 19(7): 409–424. DOI: 10.1038/s41579-021-00573-0.

Hodcroft EB, Maio ND, Lanfear R, et al. (2021) Want to track pandemic variants faster? Fix the bioinformatics bottleneck. Nature 591(7848). 7848. Nature Publishing Group: 30–33. DOI: 10.1038/d41586-021-00525-x.

Hon C-C, Lam T-Y, Shi Z-L, et al. (2008) Evidence of the Recombinant Origin of a Bat Severe Acute Respiratory Syndrome (SARS)-Like Coronavirus and Its Implications on the Direct Ancestor of SARS Coronavirus. Journal of Virology 82(4). American Society for Microbiology: 1819–1826. DOI: 10.1128/JVI.01926-07.

Jackson B, Boni MF, Bull MJ, et al. (2021) Generation and transmission of interlineage recombinants in the SARS-CoV-2 pandemic. Cell. DOI: 10.1016/j.cell.2021.08.014.

Katoh K and Standley DM (2013) MAFFT Multiple Sequence Alignment Software Version 7: Improvements in Performance and Usability. Molecular Biology and Evolution 30(4): 772–780. DOI: 10.1093/molbev/mst010.

Koelle DM, Norberg P, Fitzgibbon MP, et al. (2017) Worldwide circulation of HSV-2 × HSV-1 recombinant strains. Scientific Reports 7(1): 44084. DOI: 10.1038/srep44084.

Komissarov AB, Safina KR, Garushyants SK, et al. (2021) Genomic epidemiology of the early stages of the SARS-CoV-2 outbreak in Russia. Nature Communications 12(1). 1. Nature Publishing Group: 649. DOI: 10.1038/s41467-020-20880-z.

Korber B, Fischer WM, Gnanakaran S, et al. (2020) Tracking Changes in SARS-CoV-2 Spike: Evidence that D614G Increases Infectivity of the COVID-19 Virus. Cell 182(4): 812–827.e19. DOI: 10.1016/j.cell.2020.06.043.

Lai A, Bergna A, Acciarri C, et al. (2020) Early phylogenetic estimate of the effective reproduction number of SARS-CoV-2. Journal of Medical Virology 92(6): 675–679. DOI: 10.1002/jmv.25723.

Lam TT-Y, Hon C-C and Tang JW (2010) Use of phylogenetics in the molecular epidemiology and evolutionary studies of viral infections. Critical Reviews in Clinical Laboratory Sciences 47(1): 5–49. DOI: 10.3109/10408361003633318.

Lam-Hine T, McCurdy SA, Santora L, et al. (2021) Outbreak Associated with SARS-CoV-2 B.1.617.2 (Delta) Variant in an Elementary School – Marin County, California, May-June 2021. MMWR. Morbidity and Mortality Weekly Report 70(35): 1214–1219. DOI: 10.15585/mmwr.mm7035e2.

Lo SW and Jamrozy D (2020) Genomics and epidemiological surveillance. Nature Reviews Microbiology 18(9): 478–478. DOI: 10.1038/s41579-020-0421-0.

Martin D and Rybicki E (2000) RDP: detection of recombination amongst aligned sequences. Bioinformatics (Oxford, England) 16(6): 562–563. DOI: 10.1093/bioinformatics/16.6.562.

Martin DP, Murrell B, Golden M, et al. (2015) RDP4: Detection and analysis of recombination patterns in virus genomes. Virus Evolution 1(1). DOI: 10.1093/ve/vev003.

Martin DP, Varsani A, Roumagnac P, et al. (2021) RDP5: a computer program for analyzing recombination in, and removing signals of recombination from, nucleotide sequence datasets. Virus Evolution 7(1). DOI: 10.1093/ve/veaa087.

Matsen FA, Kodner RB and Armbrust EV (2010) pplacer: linear time maximum-likelihood and Bayesian phylogenetic placement of sequences onto a fixed reference tree. BMC Bioinformatics 11(1): 538. DOI: 10.1186/1471-2105-11-538.

Maxmen A (2021) One million coronavirus sequences: popular genome site hits mega milestone. Nature 593(7857). 7857. Nature Publishing Group: 21–21. DOI: 10.1038/d41586-021-01069-w.

McBroome J, Thornlow B, Hinrichs AS, et al. (2021) A Daily-Updated Database and Tools for Comprehensive SARS-CoV-2 Mutation-Annotated Trees. Molecular Biology and Evolution (msab264). DOI: 10.1093/molbev/msab264.

Minh BQ, Schmidt HA, Chernomor O, et al. (2020) IQ-TREE 2: New Models and Efficient Methods for Phylogenetic Inference in the Genomic Era. Molecular Biology and Evolution 37(5): 1530–1534. DOI: 10.1093/molbev/msaa015.

Nicholls SM, Poplawski R, Bull MJ, et al. (2020) MAJORA: Continuous integration supporting decentralised sequencing for SARS-CoV-2 genomic surveillance. bioRxiv. Cold Spring Harbor Laboratory: 2020.10.06.328328. DOI: 10.1101/2020.10.06.328328.

Oude Munnink BB, Nieuwenhuijse DF, Stein M, et al. (2020) Rapid SARS-CoV-2 whole-genome sequencing and analysis for informed public health decision-making in the Netherlands. Nature Medicine 26(9): 1405–1410. DOI: 10.1038/s41591-020-0997-y.

Page RDM and Holmes EC (1998) Molecular Evolution: A Phylogenetic Approach. Oxford; Malden, MA: Blackwell Science.

Price MN, Dehal PS and Arkin AP (2010) FastTree 2 – Approximately Maximum-Likelihood Trees for Large Alignments. PLOS ONE 5(3). Public Library of Science: e9490. DOI: 10.1371/journal.pone.0009490.

Rambaut A, Holmes EC, O’Toole Á, et al. (2020) A dynamic nomenclature proposal for SARS-CoV-2 lineages to assist genomic epidemiology. Nature Microbiology 5(11). 11. Nature Publishing Group: 1403–1407. DOI: 10.1038/s41564-020-0770-5.

Richard D, Shaw LP, Lanfear R, et al. (2021) A phylogeny-based metric for estimating changes in transmissibility from recurrent mutations in SARS-CoV-2. preprint, 7 May. Genomics. DOI: 10.1101/2021.05.06.442903.

Sankoff D (1975) Minimal Mutation Trees of Sequences. SIAM Journal on Applied Mathematics 28(1). Society for Industrial and Applied Mathematics: 35–42. DOI: 10.1137/0128004.

Shu Y and McCauley J (2017) GISAID: Global initiative on sharing all influenza data – from vision to reality. Eurosurveillance 22(13). European Centre for Disease Prevention and Control: 30494. DOI: 10.2807/1560-7917.ES.2017.22.13.30494.

Simon-Loriere E and Holmes EC (2011) Why do RNA viruses recombine? Nature Reviews Microbiology 9(8): 617–626. DOI: 10.1038/nrmicro2614.

Tange O (2011) GNU Parallel - The Command-Line Power Tool.;login: The USENIX Magazine 36(1): 42–47. DOI: http://dx.doi.org/10.5281/zenodo.16303.

Turakhia Y, Thornlow B, Hinrichs A, et al. (2021) Pandemic-Scale Phylogenomics Reveals Elevated Recombination Rates in the SARS-CoV-2 Spike Region. 5 August. DOI: 10.1101/2021.08.04.455157.

Turakhia Y, Thornlow B, Hinrichs AS, et al. (2021) Ultrafast Sample placement on Existing tRees (UShER) enables real-time phylogenetics for the SARS-CoV-2 pandemic. Nature Genetics 53(6): 809–816. DOI: 10.1038/s41588-021-00862-7.

Volz E, Mishra S, Chand M, et al. (2021) Transmission of SARS-CoV-2 Lineage B.1.1.7 in England: Insights from linking epidemiological and genetic data. preprint, 4 January. Infectious Diseases (except HIV/AIDS). DOI: 10.1101/2020.12.30.20249034.

Wu F, Zhao S, Yu B, et al. (2020) A new coronavirus associated with human respiratory disease in China. Nature 579(7798): 265–269. DOI: 10.1038/s41586-020-2008-3.

